# 3D-printed moulds for image-guided surgical biopsies: an open source computational platform

**DOI:** 10.1101/658831

**Authors:** Mireia Crispin-Ortuzar, Marcel Gehrung, Stephan Ursprung, Andrew B Gill, Anne Y Warren, Lucian Beer, Ferdia A Gallagher, Thomas J Mitchell, Iosif A Mendichovszky, Andrew N Priest, Grant D Stewart, Evis Sala, Florian Markowetz

**Author notes:** These authors contributed equally to this work (Mireia Crispin-Ortuzar, Marcel Gehrung, Stephan Ursprung). Shared senior authorship (Grant D Stewart, Evis Sala, Florian Markowetz).

## Abstract

**PURPOSE:** Spatial heterogeneity of tumours is a major challenge in precision oncology. The relationship between molecular and imaging heterogeneity is still poorly understood, as it relies on the accurate co-registration of medical images and tissue biopsies. tumour moulds can guide the localization of biopsies, but their creation is time consuming, technologically challenging, and difficult to interface with routine clinical practice. These hurdles have so far hindered the progress in the area of multiscale integration of tumour heterogeneity data.

**METHODS:** We have developed an open source computational framework to automatically produce patient-specific 3D-printed moulds that can be used in the clinical setting. Our approach achieves accurate co-registration of sampling location between tissue and imaging, and integrates seamlessly with clinical, imaging and pathology workflows.

**RESULTS:** We applied our framework to patients with renal cancer undergoing radical nephrectomy. We created personalised moulds for five patients, obtaining Dice similarity coefficients between imaging and tissue sections ranging from 0.86 to 0.93 for tumour regions, and between 0.70 and 0.76 for healthy kidney. The framework required minimal manual intervention, producing the final mould design in just minutes, while automatically taking into account clinical considerations such as a preference for specific cutting planes.

**CONCLUSION:** Our work provides a robust and automated interface between imaging and tissue samples, enabling the development of clinical studies to probe tumour heterogeneity on multiple spatial scales.

## INTRODUCTION

Molecular tumour profiling is used to stratify patients and identify new actionable targets for precision therapeutics. The assessment is typically based on data from a single tumour biopsy^1^. Often, however, tumours display such a high degree of heterogeneity that a single tissue sample is insufficient to capture the full molecular landscape of the disease^2^. A prime example of such spatial heterogeneity is renal cell carcinoma (RCC), which has been shown to be radiologically, genetically, and metabolically heterogeneous^3–5^. Macroscopic regions with distinct genotypes can be identified within a single tumour through multi-regional sampling^3,6^. In parallel, radiological imaging provides non-invasive, three-dimensional information on phenotypic heterogeneity^7,8^. The fact that RCC displays spatial heterogeneity at such disparate physical scales suggests that a combined approach to integrate the relevant data sources (i.e. genomics, transcriptomics, radiomics) is needed to unravel the complexity of the disease^9^ and the genomic evolution of the tumour^4,10–12^. The foundation of a combined analysis is the accurate spatial co-registration of imaging data and biopsies. However, typically multi-regional tumour biopsies are obtained after nephrectomy, when image guidance is no longer possible. The challenge of co-registering *in vivo* images to resected tumours has been addressed in other contexts. Previous solutions included holding the specimen with a cradle^13^ or solidified agar^14^. However, these approaches had several disadvantages, including not being clinically usable, or not providing accurate orientation. More recently, personalised 3D moulds have been used to improve the accuracy of co-registration in prostate cancer^15–17^ and ovarian cancer studies^18^.

In RCC, however, 3D-printed moulds remain comparatively underexplored^19^, as the disease presents unique challenges. The first challenge arises from the pathology guidelines for assessment of radical nephrectomy specimens, which require optimal visualisation of the renal sinus–tumour interface. The most commonly adopted initial plane of incision is along the long axis at midpoint, with further sectioning usually perpendicular to this plane^20–22^. Thus, the sectioning planes are in general not the same as those used for imaging. An additional challenge is that pathologists need to preserve the integrity of structures, which are required for staging, such as the renal vein. Finally, the specimen is often covered by perinephric fat^23^, which further complicates the procedure and can make it impossible to identify relevant structures. Because of these restrictions, previous 3D-printing-based co-registration methods for RCC have either been limited to pre-clinical models^24^, or have only focused on early-stage partial nephrectomy cases^25^, where the fat-free resection margin can be used as a base. In addition, none of the preivous methods addressed the issue of having different sectioning and imaging planes. New methods are therefore needed to accurately match macroscopic habitats defined by imaging to specific tissue regions. Importantly, these methods need to integrate smoothly into the clinical pathway to allow future use in clinical trials and potentially clinical practice.

Here we report the design and implementation of an open-source computational framework to create image-based patient-specific tumour moulds. The moulds enable the co-registration of surgical tissue samples to pre-surgical multiparametric magnetic resonance imaging (MRI) in patients undergoing radical nephrectomy for suspected RCC. Our methodology is fully automated, producing ready-to-print 3D designs directly from the MRI segmentation. It is also tailored for seamless integration with the clinical workflow. In particular, it can deal with any desired sectioning plane, and is based on a robust landmark system that ensures accurate co-registration even in specimens obscured by a thick adipose layer. Although the framework was designed for renal cancer, it could be easily adapted to any other type of solid tumour. As such, it constitutes a substantial step forward towards streamlining the creation of datasets with accurately matched imaging, histological and genomics data. Below we present the computational details of the framework and validate its performance on five radical nephrectomy cases.

## RESULTS

### Key concepts

We are presenting a framework to create moulds that can assist the tumour sampling process by co-registering tumour sections with MRI slices. The mould is a three-dimensional block, with vertical slots that guide the sectioning, and a cavity designed to precisely fit the resected specimen (Figure 1a). The shape of the cavity is derived from the regions of interest drawn by a radiologist on a MRI scan. The 3D modelling process involves several steps, including volume creation, re-orientation, smoothing, mesh creation, and the addition of slots and guides (Figure 1a). All steps proceed automatically, and they integrate with the clinical workflow (Figure 1b). The code is available online on doi:10.5281/zenodo.3066304.

**FIG. 1.**
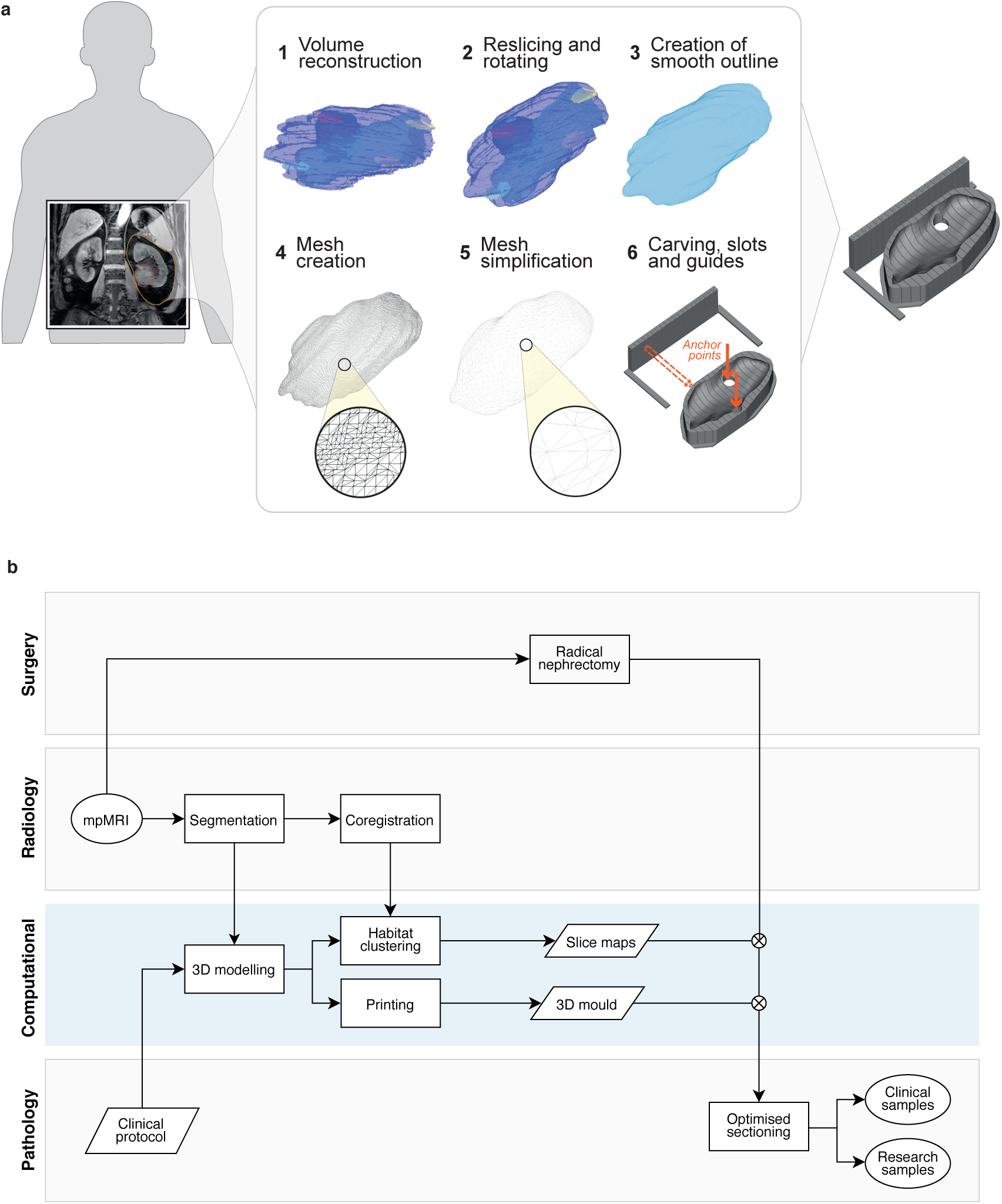
A computational framework to create image-based patient-specific tumour moulds. **(a)** The schematic depicts the various steps of the method, bridging from MRI scans to spatially targeted surgical biopsies. The method starts with the delineation of a MRI scan, which is then re-oriented, carved out of a 3D-printed mould, and used for spatially accurate surgical biopsies. The slots of the mould guide the knife for cutting. **(b)** Flow chart of the different analysis steps performed by the radiology, surgery, pathology and computational groups to ensure seamless integration between the clinical and research arms. The blue box highlights the computational steps of the pipeline.

### Automated 3D modelling

#### Step 1: Image segmentation

Our approach requires two types of regions of interest (ROIs) to be drawn on the images: tissue segmentations and anatomic landmarks. Tissue segmentations are needed to define the mould cavity and to test the spatial accuracy of the framework. They include the tumour, normal kidney, and perinephric fat. Combined, they form the global outline of the specimen, which defines the shape of the mould.

In addition, at least four anatomic landmarks are needed to determine the correct orientation of the specimen inside the mould. The first two are the upper and lower poles of the kidney, which ensure that the kidney can be sectioned along or transverse to its long axis at midpoint^20^. The other two anatomic landmarks are the hilum (exit point of renal vessels and ureter) and the points in the tumour and/or normal kidney with the thinnest fat coverage, referred to as ‘contact points’. They are used to ensure that the specimen is accurately positioned.

#### Step 2: Image orientation

Our approach controls the orientation of the specimen within the mould. The first orientation challenge concerns the direction along which the specimen has to be sectioned, following pathology protocols for renal cancer staging. To address this, we apply a 3D rotation to the images and create new slices that align with the preferred sectioning plane, which is defined by the tumour centroid and the upper and lower poles.

The second challenge concerns the need to accurately orient the specimen in the mould, even when it is covered in perinephric fat. We overcome this challenge by defining reference landmarks that are expected to be exposed and identifiable in the specimen, and placing them at the base of the mould. These points act as anchors that ensure that the specimen is correctly positioned. The points are marked in the mould by carving 2 cm holes in the base of the mould that enable the pathologist to see and feel them (red arrows in Figure 1a). The two landmark points used for this purpose are the hilum and the tumour contact point.

Once the image has been rotated, we extract the outline volume needed for the mould and smooth the surface using a Gaussian kernel. The final output is a three-dimensional integer matrix that embeds the correctly oriented volume as well as the location of the landmark points. This part of the process is implemented in MATLAB.

#### Step 3: Mould generation and 3D printing

The mould generation process consists of several steps (Figure 1a). First, the volumetric matrix obtained previously is converted into a mesh, and then simplified by face reduction, adaptive remeshing, Laplacian smoothing and Taubin smoothing.

Once the mesh is smooth enough for printing, it is carved off from a solid block-shaped base, and vertical slots are created to guide the knife during sectioning. In addition, a set of vertical guides is added to one side of the mould, to aid with the positioning of the knife. The location of the inter-slot spaces in both the guides and the mould is designed to match the exact location of the imaging slices of interest. Additionally, the guides are numbered such that particular slices can easily be identified and compared to imaging. Finally, we carve the reference holes at the bottom of the mould with a diameter of 2 cm at the hilum and contact landmark points. This part of the process is implemented in Slic3r (Prusa Research, Czech).

### Validation in a pilot study

The methodology was validated using specimens from five patients with renal tumours who underwent radical nephrectomy (Figure 2). Relevant clinical data can be found in Table I, and additional information can be found in the Methods section.

**TABLE I.** MRI Parameters

| Case number | Gender | Age | Tumour type | Tumour stage | Nodal stage | Tumour grade | Time between imaging and surgery [days] | Tumour volume [ml] |
| --- | --- | --- | --- | --- | --- | --- | --- | --- |
| 1 | M | 70 | Clear cell RCC | pT3a | pNx | 3 | 14 | 145 |
| 2 | M | 81 | Clear cell RCC | pT3a | pNx | 4 | 29 | 218 |
| 3 | F | 60 | Clear cell RCC | pT3a | pNx | 2 | 19 | 89 |
| 4 | M | 51 | Clear cell RCC | pT3a | pN0 | 4 | 22 | 107 |
| 5 | M | 47 | Rhabdomyosarcoma | pT4 | pN0 | ungraded | 11 | 1342 |

M: Male, F: Female, RCC: Renal cell carcinoma

**FIG. 2.**
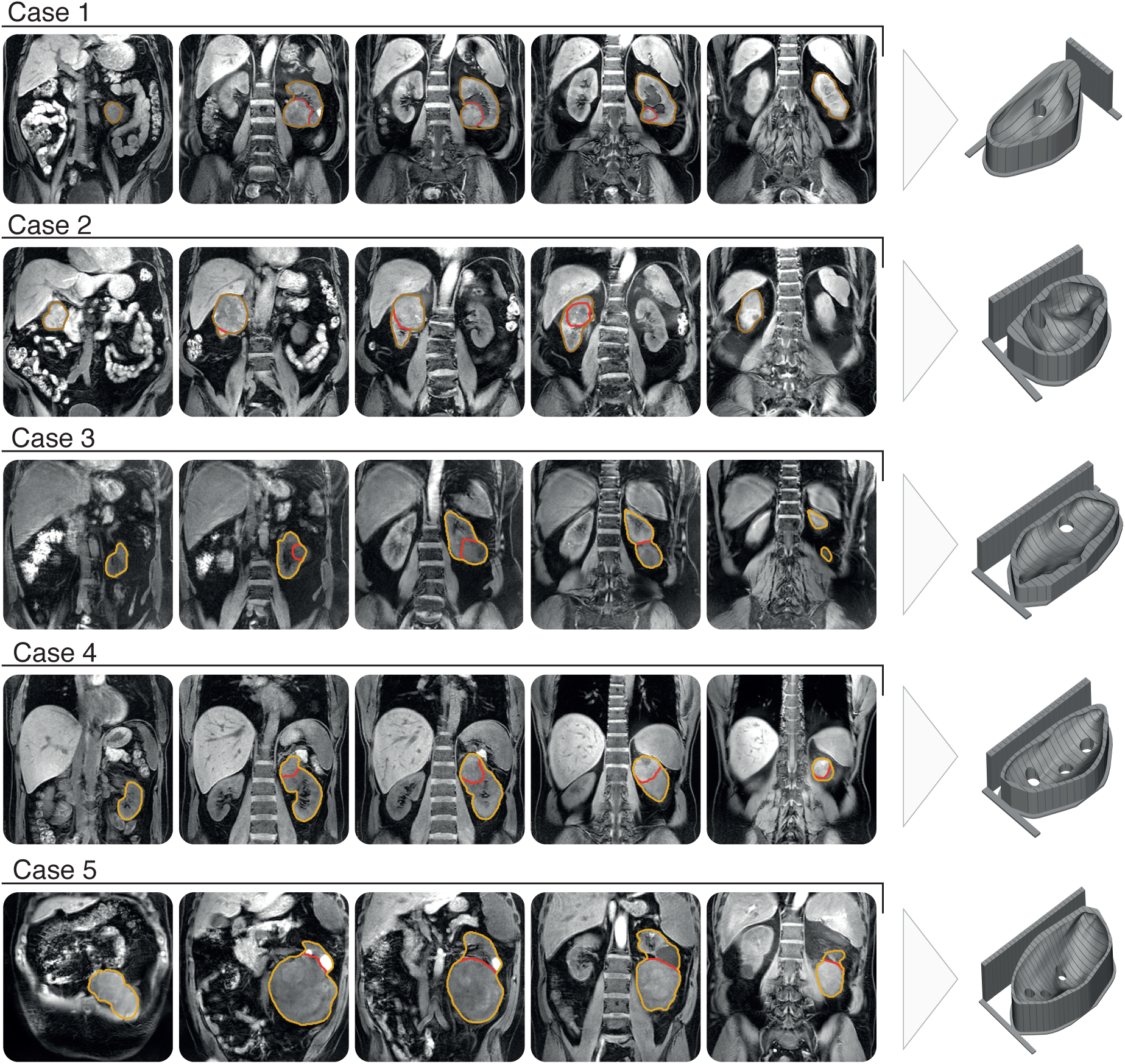
Optimised, patient-specific tumour moulds. Representative T1w MRI slices and corresponding 3D renderings of the tumour moulds created for the five cases included in the study.

tumour, normal kidney and perinephric fat were segmented manually on a pre-surgical T1w MRI image, as well as the hilum, tumour and kidney contact point and kidney poles. For the first patient, the renal pelvis was also segmented. The segmentations were checked by a radiologist with 15 years of experience in genitourinary imaging (ES).

We generated and 3D-printed moulds for each patient using the computational framework described above. After discussion with the pathologist, it was decided that the first case would be sectioned longitudinally to the kidney, while the other four were sectioned transversally.

The automated design and generation of each mould took less than 5 minutes per patient. Manual verification of the segmentation and mould results took between 10 and 20 minutes. Printing each mould took between 12 and 24 hours.

The specimens were placed in the mould and sectioned 20 minutes after nephrectomy. The resection margins were inked for R-staging and all the perinephric fat was preserved. A slice where all the habitats of the tumour were present, as well as being sufficiently separated from the hilum, was chosen for sectioning in each case. Cuts were made with a 12-inch CellPath Brain Knife.

#### Anatomical landmark validation

Based on these data we validated the anatomical landmarks and the functional signal. In the first case, the selected slice resulted in a clean longitudinal cut of the kidney, including the renal pelvis, and a cross-section of the tumour, as illustrated in Figure 3. The tumour presented two hemorrhagic areas and a necrotic core. The other four cases were sectioned transversally, with cases 2 and 3 including large portions of normal kidney.

**FIG. 3.**
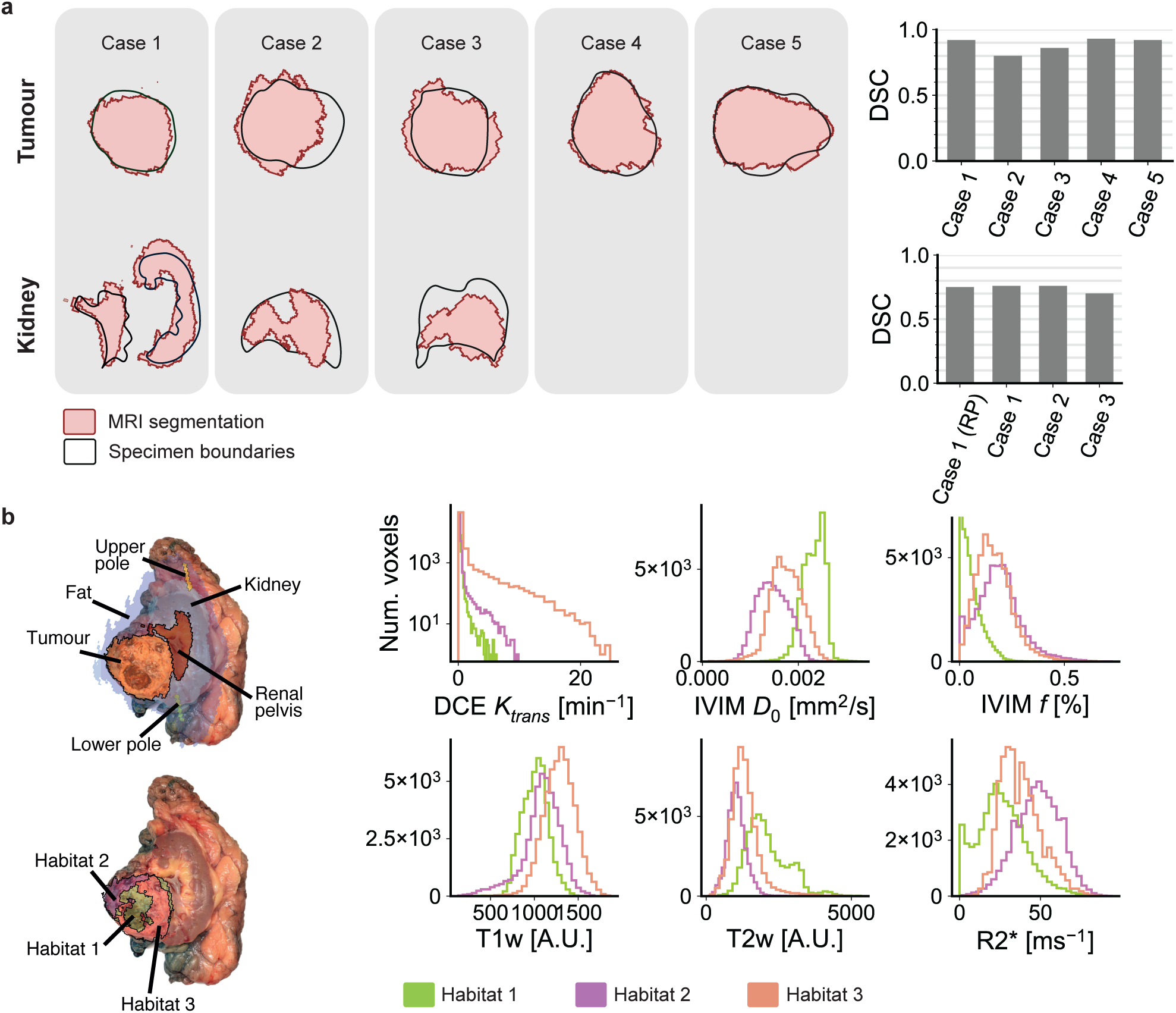
Validation results. (a) Overlay of the tissue region boundaries (black) and the corresponding MRI segmentations (red) for tumour and kidney regions. DSCs are calculated for tumour and kidney tissues separately. RP indicates the renal pelvis. (b) Left: Overlay of a photograph of the section from the first case and the corresponding MRI maps, including anatomical region segmentations (top) and multi-parametric tumour habitats (bottom). Right: Relative distributions of imaging parameters for the three tumour habitats.

For each case, the slice was placed on a flat surface and photographed. We then manually contoured the reference tissues (tumour, kidney, and renal pelvis where visible) on the tissue photograph. We co-registered the MRI segmentations and tissue contours manually, obtaining Dice Similarity Coefficients (DSCs)^26^ of 0.92, 0.80, 0.86, 0.93 and 0.92 respectively for the five tumour ROIs, as shown in Figure 3a. The three regions of interest containing healthy kidney yielded DSCs of 0.76, 0.76 and 0.70 respectively. For the first case, the renal pelvis yielded a DSC of 0.75.

#### Functional signal validation

Motivated by the presence of a necrotic core in the first patient, we performed a further validation step based on the spatial distribution of different functional imaging parameters inside the tumour. Multiparametric MRI images were co-registered and used to define spatial habitats using *k*-means clustering. In particular, we used T1w and T2w images, T1 map, *K*^trans^ from dynamic contrast enhanced (DCE) MRI as a measure of tumour vascular leakage, the D0 diffusion coefficient and perfusion fraction from IVIM MRI imaging (*f*) as a measure of cellularity and tumour perfusion, and R2*, as a measure of oxygenation. We found three distinct habitats, as shown in Figure 3b.

All three habitats presented with distinct distributions with respect to perfusion fraction *f, K*^trans^ and R2* maps, as shown in Figure 3b. We found habitat 1 to be poorly perfused and have a high diffusivity, T1w hypointensity and T2w hyperintensity. This habitat overlapped with the necrotic area found in the resected specimen.

Habitats 2 and 3 showed similar parametric distributions. Habitat 2 was adjacent to the kidney and showed the highest levels of *K*^trans^. Habitat 3 showed the lowest diffusion levels, as well as high R2*.

## DISCUSSION

Capturing the full complexity of the disease is very challenging in cases like RCC, where tumours typically display a high degree of spatial heterogeneity both at the imaging and genomics level. In this paper we have presented a new computational framework that overcomes a key challenge for the combined analysis of imaging and genomics data: the need to accurately match macroscopic habitats defined by imaging to specific tissue regions in an automated way and without disrupting routine clinical practices. By integrating smoothly into clinical practice, our methodology has the potential to be widely applicable in clinical trials and therefore enable the creation of unprecedented datasets with matched imaging, histological and genomics data.

### An open-source automated platform for mould creation

Our framework successfully integrated all the steps to automatically produce 3D-printable moulds directly from MRI segmentations. This facilitates the inclusion of mould-guided samples into clinical studies, because moulds can be generated fast with minimal additional workload.

### Mapping imaging and sectioning planes

Our approach was designed to address one of the limitations of previous 3D-printing-based co-registration methods, which assumed that tumours can be sectioned along the same plane that was used for MRI imaging. This assumption generally interferes with pathology protocols. Commandeur *et al.* proposed a methodology to co-register histological planes to MRI slices for prostate cancer^27^. However, this co-registration has to be performed *a posteriori* and therefore the surgical biopsies would need to be obtained without image guidance, which might result in sub-optimal tumour sampling^10^. Instead, our approach uses a landmark system based on the definition of two reference points drawn by the radiologist on the MRI scan (the upper and lower poles of the kidney). These points are then used to define the rotation to be applied to the images. We found that the rotation successfully provided the expected longitudinal or transversal cuts of the kidney.

### Accurate co-registration in the presence of perinephric fat

The second challenge addressed by our approach is the presence of perinephric fat, which adds two complications to the tissue co-registration process: the difficulty in predicting the exact shape of the resected specimen, as the definition of optimal margins is controversial^28^; and the lack of an anatomical frame of reference to correctly position the specimen in the mould. Removing or trimming the fat may interfere with clinical practice, as it could compromise the surgical margins, which need to be evaluated for the presence of tumour cells^29^. A solution has been previously proposed for partial nephrectomy cases, using the inner parenchymal surface of the tumour as the base of the mould^25^. This method involved the surgeon inserting fiducial markers into the tumour during surgery, which interrupts the routine clinical pathway. In addition, partial nephrectomy is only recommended to treat small renal masses^30^, so more advanced cases, which have typically poorer outcomes and are therefore of particular clinical relevance^31^ would not be tractable with this approach.

Our methodology instead relies on a second set of key landmarks that can be used to orient the specimen even when there is a large component of fat. These reference points are placed at the base of the mould and marked with holes that allow the pathologist to confirm their correct positioning. This approach resulted in an accurate co-registration between imaging and resected specimen in five specimens corresponding to renal cancers of stages 3 and 4. In particular, we found that anatomical image segmentations agreed with the corresponding tissue outlines after mould-assisted sectioning, with DSCs ranging between 0.86 and 0.93 for tumour regions, and between 0.70 and 0.76 for healthy kidney regions.

In addition, we observed that the tumour habitats identified from multiparametric MRI images from case number 1 coincided with observable features of the tissue. In particular, habitat 1 presented all the characteristics of necrotic tissue (poor perfusion, high diffusion, T1w hypointensity and T2w hyperintensity), and indeed coincided with the necrotic core of the tumour^32^. Similarly, habitat 3, which was closest to the normal kidney and therefore potentially could have better vascular access, was found to have high *K*^trans^.

As expected, there was a thick layer of fat surrounding the specimens, which made it difficult to see the kidney or identify its orientation by simple visual inspection. This would have been a challenge even in the standard clinical setting, but the mould generally provided useful support and assistance.

### Limitations of the approach

Our approach shares some limitations with most other co-registration approaches. First of all, there is a time constraint between imaging and surgery. In this study imaging occurred between 2 and 4 weeks before surgery, which could have resulted in anatomical changes and therefore a suboptimal mould design. However, typical tumour doubling times for renal cancer are large, and suggest that the effects should be minor^33,34^. Shape-wise, additional uncertainty may arise from the segmentation of the structures on the MRI images. Although several approaches for semi-automatic segmentation of kidney tumours exist^35–37^, the preferred option is still manual contouring. Our methodology requires the additional delineation of perinephric fat, for which manual contouring, after discussion with the surgeon, is preferred. Although placing the point with the least fat coverage at the bottom of the mould helps reduce the uncertainty, intra-operative decisions may result in a different fat distribution. Having a single-sided mould (without an upper half) means that changes in the upper side of the specimen do not impact the accuracy, but any variations in the other half might do.

### Impact and future work

The methodology we have presented here will be a core element of the WIRE renal cancer trial^38^. By tightly integrating into the workflows of clinical trials, our methodology will enable the creation of large spatially-matched multiscale datasets including radiomics, genomics and histology data.

## MATERIALS AND METHODS

### Code

All the code necessary to reproduce these results, including volume orientation, 3D mould design, 3D printing, and habitat generation, can be found in doi:10.5281/zenodo.3066304.

### Ethics and patient cohort

The method was designed as part of a physiological study currently being undertaken at the University of Cambridge with the aim of integrating of imaging and tissue based biomarkers to unravel tumour heterogeneity in renal cancer. Informed consent was obtained for the Molecular Imaging and Spectroscopy with Stable Isotopes in Oncology and Neurology - substudy in renal cancer (MISSION) after prior approval by the East of England - Cambridge South ethics committee (REC: 15/EE/0378).

All patients recruited for the MISSION study between December 2018 and December 2019, a total of 8, were considered for 3D mould printing. Of those, three were excluded from the validation study presented here: one due to having withdrawn consent to imaging (no mould was designed), one due to having a paraganglioma (no mould was designed), and one due to not having tissue photographs for anatomical validation. The analysis presented here is based on the remaining five patients.

### MRI data acquisition

The 3D model of the tumour was designed based on a T1-weighted (T1w) MRI scan acquired using a Dixon imaging sequence (Table II) acquired between two and four weeks before surgery on a clinical 3T MRI (Discovery MR750, GE Healthcare, Waukesha, WI). Regions of interest (ROIs) were manually delineated by a radiologist (SU, with 2 years of experience in genitourinary imaging) on each slice of the MRI scan using OsiriX (Version 10.0.0^39^). The contours were drawn on coronal unenhanced T1w images using registered T2w and post-contrast T1w images to verify the accuracy of the ROIs. The segmentation was independently reviewed by a second radiologist (ES, with 15 years of experience in genitourinary imaging). ROIs were exported from OsiriX to comma separated value files (.csv) encoding the coordinates of the edges of the ROI on each slice using the Export ROIs plugin (Version 1.9). The centroid of each ROI was calculated as the mean of all *x, y* and *z* coordinates of the voxels within it.

**TABLE II.**
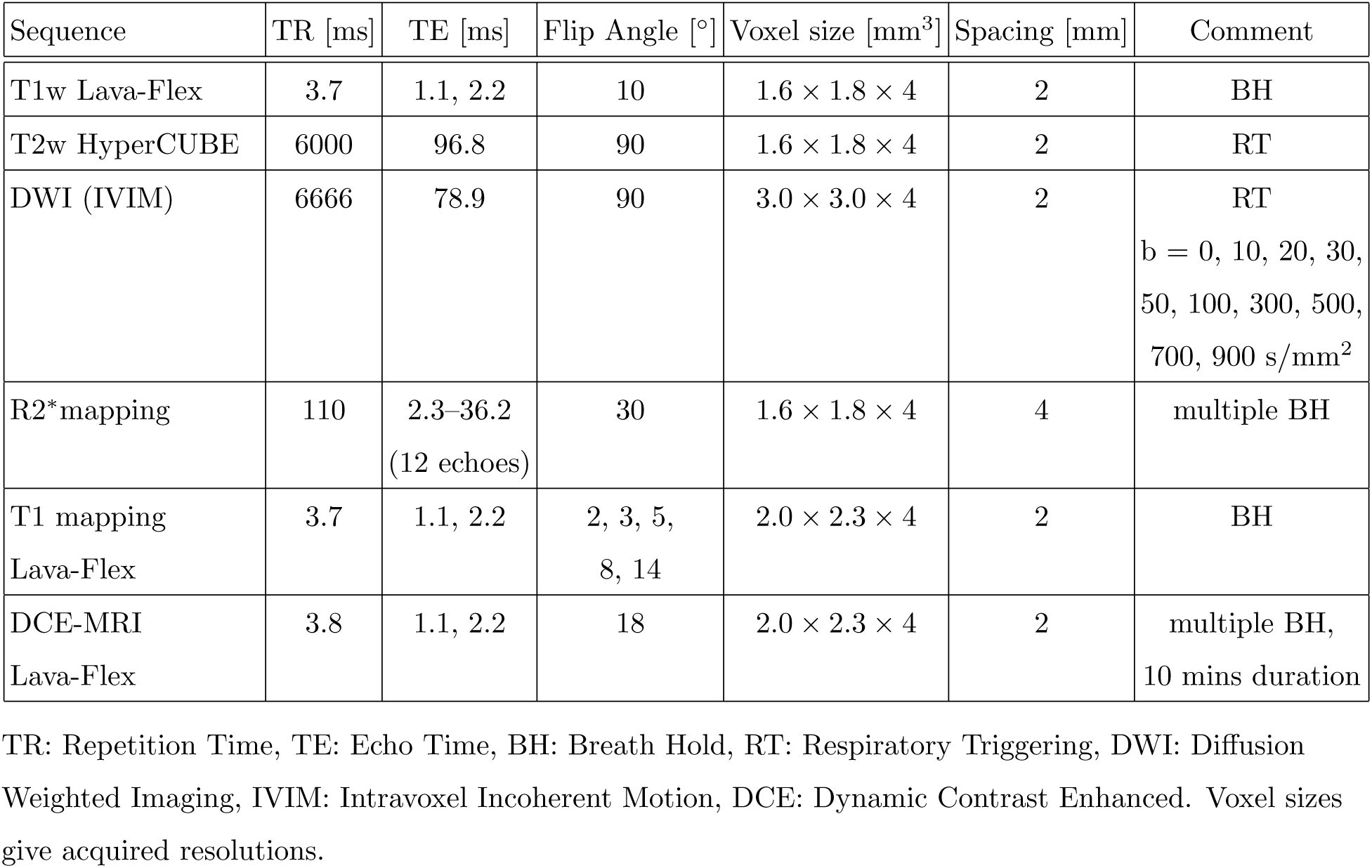
MRI Parameters

| Sequence | TR [ms] | TE [ms] | Flip Angle [°] | Voxel size [mm<sup>3</sup>] | Spacing [mm] | Comment |
| --- | --- | --- | --- | --- | --- | --- |
| T1w Lava-Flex | 3.7 | 1.1, 2.2 | 10 | $1.6 \times 1.8 \times 4$ | 2 | BH |
| T2w HyperCUBE | 6000 | 96.8 | 90 | $1.6 \times 1.8 \times 4$ | 2 | RT |
| DWI (IVIM) | 6666 | 78.9 | 90 | $3.0 \times 3.0 \times 4$ | 2 | RT<br>b = 0, 10, 20, 30,<br>50, 100, 300, 500,<br>700, 900 s/mm<sup>2</sup> |
| R2* mapping | 110 | 2.3–36.2<br>(12 echoes) | 30 | $1.6 \times 1.8 \times 4$ | 4 | multiple BH |
| T1 mapping<br>Lava-Flex | 3.7 | 1.1, 2.2 | 2, 3, 5,<br>8, 14 | $2.0 \times 2.3 \times 4$ | 2 | BH |
| DCE-MRI<br>Lava-Flex | 3.8 | 1.1, 2.2 | 18 | $2.0 \times 2.3 \times 4$ | 2 | multiple BH,<br>10 mins duration |

TR: Repetition Time, TE: Echo Time, BH: Breath Hold, RT: Respiratory Triggering, DWI: Diffusion Weighted Imaging, IVIM: Intravoxel Incoherent Motion, DCE: Dynamic Contrast Enhanced. Voxel sizes give acquired resolutions.

### Image pre-processing

Before generation of parameter maps, deformable motion correction was applied in MATLAB (Mathworks, Natick, MA) and utilizing ANTs/ITK^40^. In the case of DWI-MRI this was applied across acquisitions with differing b-values; in the case of DCE-MRI, this was applied across acquisition time-points and the associated T1 maps were transformed accordingly. Parameter maps were then generated using MATLAB in the case of DWI-IVIM, and using MIStar (Apollo Medical Imaging Technology, Melbourne, Australia) in the case of DCE-MRI, employing the Tofts model^41^ and a model arterial input function. R2* maps were generated at source on the MRI scanner using standard manufacturer software. All parameter map volumes were then aligned to the T1-weighted reference series used to prepare the mould. This was performed in two stages: first each parameter map volume was resampled into the space of the T1w reference series. Finally, and only if necessary, a rigid registration transform to more closely align the map with the reference image was determined manually using the software package ITK-SNAP; this transform was then applied to the parameter map volume using MATLAB.

### Mould orientation

The method proceeds as follows. First, the MRI scan is re-sampled to achieve an isotropic resolution of 1 × 1 × 1 mm^3^ using nearest neighbour interpolation, as implemented in CERR^42^, an open source MATLAB environment for radiology research. Then, two three-dimensional rotations are applied. Several vectors connecting the structure centroids are defined to guide the re-orientation process, as follows:

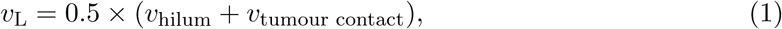

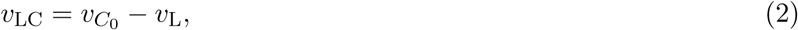

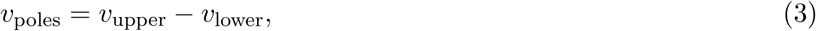

where *v*_*i*_ indicates the coordinates of the centroid of structure *i*, with *v*_upper_ representing the centroid of the upper pole, and *v*_lower_ the centroid of the lower pole. The first rotation aligns *v*_poles_ with the *y* axis. The second rotation is performed around the *y* axis, aligning the *x* − *z* projection of *v*_LC_ with the *z* axis. Combined, the two rotations ensure that the orientation conditions are satisfied. Other rotation choices could also be easily implemented.

Before extracting and exporting the re-oriented volume for mould design, the surface is smoothed using 3D Gaussian filtering with a convolution kernel of size 9 × 9 × 9 voxels and standard deviation of 3 voxels. Finally, the MRI images are sliced along the *x* − *z* plane with a spacing of 1 cm. These are used to build reference maps that will later guide the tissue sampling process; they also coincide with the location of the mould’s slots.

### Design optimisation & mould generation

The resliced tumour segmentation was exported from MATLAB and imported into a Python script for post-processing and mould generation. First, the marching cubes algorithm^43^ was applied on the 3D volume for conversion to a mesh consisting of faces and vertices. Second, in order to ensure integrity of the resulting tumour mesh close vertices were merged, duplicate faces and vertices removed, faces from non-manifold edges removed, and all face normals orientations inverted. Third, the number of faces was reduced to 5000 max. by performing quadric edge collapse decimation to simplify the mesh and reduce computational overhead. Fourth, the first smoothing step with a Laplacian kernel was performed. Fifth, as Laplacian smoothing can result in geometric issues in certain scenarios, faces were again removed from non-manifold edges as well as duplicated faces and vertices removed. Sixth, Taubin smoothing was performed to remove remaining irregularities. Last, remaining holes in the mesh were closed to ensure a continuous surface for mould generation and printing. Detailed parameters for each step can be found in the file *filter.mlx* on doi:10.5281/zenodo.3066304.

### 3D printing

The model was sliced using PrusaSlicer (Prusa Research, Czech) and printed with 0.2 mm layer height on a Prusa i3 MK3 printer loaded with RS PRO PLA filament (RS Components, UK).

### Habitat clustering

In order to guide the process of tissue sampling, imaging maps were created for each tumour slice. The maps were obtained by combining multiparametric MRI images and clustering them into several spatial clusters.

Along with the reference T1w images, additional sequences were acquired to define phenotypic habitats in the first patient. In particular, the images used for clustering were the T1w and T2w images, T1 map, *K*^trans^ from DCE MRI, the diffusion coefficient and perfusion fraction from IVIM MRI imaging (*f*), and R2*. Images were obtained on a 3T MRI scanner, in coronal orientation with a slice thickness of 4 mm. Scans were corrected for motion artefacts and co-registered using rigid transformations. Additional details on the images, parameter maps, and methods can be found in Table II and the supplementary materials.

Habitats were obtained by applying *k*-means clustering on the set of co-registered images as well as the (x,y,z) coordinates corresponding to each voxel, to ensure spatial cohesion. The number of clusters was set to the maximum number that would allow taking three samples from each habitat. In practice, this translated into increasing the number of clusters until any of the habitats had an area smaller than approximately 3 cm^2^.

### Evaluation of spatial accuracy

The slice was placed on a flat surface and photographed. Tissue contours were drawn on the image, being completely blinded to the MRI segmentations. The resulting outline and the shape predicted after reorientation of the MR-segmentation were then overlayed and co-registered using manual rigid registration, maximizing the overlap between the tumour contours. The accuracy of slice position recovery was assessed post-resection by comparing the DSC of MRI segmentations and the corresponding tissue contours. This coefficient is defined as:

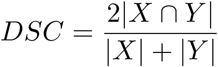

where the overlap of two binary masks X and Y (segmentations originating from different image sources) can be calcuated. The higher the DSC, the larger the overlap between the two binary masks.

## ACKNOWLEDGMENTS

The authors acknowledge the help of Gaspar Delso (GE Healthcare) and Dattesh Shanbhag (GE Global Research) for the use and ongoing support of their MRI image motion-correction programming code. MCO acknowledges support from a Borysiewicz Fellowship from the University of Cambridge and Junior Research Fellowship from Trinity College, Cambridge. MG acknowledges support from an Enrichment Fellowship from The Alan Turing Institute. This work was supported by the Wellcome Trust (095962), Cancer Research UK (CRUK; C9685/A25117, C8742/A18097, C19212/A16628, C19212/A911376, C19212/A27150, C14303/A17197, C14303/A19274), the CRUK Engineering and Physical Sciences Research Council (EPSRC) Cancer Imaging Centre in Cambridge and Manchester (C197/A16465), the Mark Foundation Institute for Integrative Cancer Medicine at the University of Cambridge, Addenbrooke’s Charitable Trust, the National Institute for Health Research (NIHR) Cambridge Biomedical Research Centre and Cambridge University Hospitals NHS Foundation Trust. Infrastructure for the Cambridge Urological Bio-repository was funded by the Cambridge Biomedical Research Campus and CRUK Cambridge Centre. Author ANP and the Human Research Tissue Bank are supported by the NIHR Cambridge Biomedical Research Centre.

